# Label propagation defines signaling networks associated with recurrently mutated cancer genes

**DOI:** 10.1101/320770

**Authors:** Merve Cakir, Sayan Mukherjee, Kris C. Wood

## Abstract

Each different tumor type has a distinct profile of genomic perturbations and each of these alterations causes unique changes to cellular homeostasis. Detailed analyses of these changes would reveal downstream effects of genomic alterations, contributing to our understanding of their roles in tumor development and progression. Across a range of tumor types, including bladder, lung, and endometrial carcinoma, we determined genes that are frequently altered in The Cancer Genome Atlas patient populations to study the effects of these alterations on signaling and regulatory pathways. To achieve this, we used a label propagation-based methodology to generate networks from gene expression signatures of mutations. Individual networks offered a comprehensive view of signaling changes represented by gene signatures, which in turn reflect the scope of molecular events that are perturbed in the presence of a given genomic alteration. Comparing different networks to each other revealed commonalities between them and biological pathways distinct genomic alterations converge on, highlighting the critical signaling events tumor dysregulate through multiple mechanisms. Finally, mutations inducing common changes to the signaling network were used to search for genomic markers of drug response, connecting shared perturbations to differential drug response.

## Introduction

Recent advances in high-throughput sequencing technologies and large-scale efforts like The Cancer Genome Atlas (TCGA) have revealed, for the first time, the landscapes of genomic alterations found within distinct tumor types, providing new insights into the mechanisms of tumor development and progression [1] [2] [3]. Further, comparisons of the patterns of genomic alterations within and across tumor types [4] [5] have revealed that tumors regulate their growth and survival through both shared and distinct mechanisms. Together, these studies underscore the importance of diverse and incompletely understood genomic changes in dictating tumors’ biological and therapeutic response characteristics.

The next step following identification of tumor genomic alterations is to understand how these alterations disturb cellular homeostasis or contribute to tumorigenesis. Connecting recurrently altered genes to the signaling or regulatory pathways that they participate in is a common starting point to achieve this goal. For instance, a common approach employed by the TCGA and related projects is to place genes into groups representative of the pathways or processes in which they are believed to function, such as MAPK signaling, PI3K signaling, and cell cycle. However, in many cases, the alteration of a cancer gene leads to cellular changes that are only partially understood and cannot be easily placed into defined pathways. Additionally, even those genomic alterations involving genes canonically implicated in a defined pathway have the potential to induce non-canonical signaling changes. Therefore, grouping genomic alterations based only on our current knowledge-based annotations can lead to an incomplete representation of their effects. Alternative approaches that enable a more comprehensive look into signaling changes can provide a more in-depth understanding of downstream effects of frequently altered cancer genes. Further, analyzing these effects across different genes may highlight common signaling events that are perturbed downstream of distinct alterations, revealing the mechanisms by which tumors achieve common outcomes (e.g., growth and survival) via distinct mechanisms.

One way of performing a detailed analysis of the effects of defined genomic alterations on signaling events is by focusing on gene expression patterns to identify genes which display dysregulated expression in the presence of a given alteration. Creating a gene expression signature by comparing mutant and wildtype samples is an established method for such an analysis. This signature, however, will often result in a sparse representation of the molecular changes associated with an alteration, as it will typically be based on strong discriminators and cannot possibly contain every gene in a dysregulated pathway. Searching for enriched Gene Ontology or functional annotation terms is a common way to better understand the molecular events represented by a given gene expression signature [6] [7]. Gene Set Enrichment Analysis (GSEA) is another commonly used method for connecting gene expression patterns to perturbations in signaling events [8]. One drawback of these enrichment based approaches, however, is the fact that they ignore connectivity within and between enriched gene sets. Thus, they yield information on the molecular events that are dysregulated but do not provide information on how the genes that contribute to these events functionally relate to one another. Additionally, these approaches lack information on the crosstalk between different gene sets that are dysregulated, making it harder to gain a unified understanding of the complete set of changes that are induced by a given genomic perturbation.

The connectivity within a given gene set, and crosstalk between different sets, offers valuable information because molecular events in a cell occur through an interconnected web of interactions. The changes that occur as a result of a given genomic alteration will spread across the molecular network through signaling cascades, rather than distinctly affecting separate sets of genes. Therefore, a more informative approach would be to create network-level views of signaling changes that are observed in the presence of a given alteration. This means that we would require an approach that can go from a gene list level to a network structure that connects these genes. There are multiple ways of achieving this. For example, one can simply connect genes to their direct interaction partners found within the gene set or introduce intermediate genes that are found along the shortest paths connecting the elements of the gene set [9] [10]. We have chosen instead to focus on another alternative, label propagation. Label propagation algorithms start with a given set of seed genes and diffuse through the network based on its specific topology to identify additional genes that are in the neighborhood of the seed genes, connecting them together. This diffusive property enables the algorithm to fully exploit the topological information offered by a given network and to discover a variety of paths that can connect a given gene set. This can create a more comprehensive representation of the starting gene set compared to linking direct neighbors or connecting them only through shortest paths. In biology, label propagation algorithms have been used to address several different problems, like predicting functions of genes based on their relationships with other well-annotated genes [11], discovering novel genes that are associated with a disease [12], [13] or differentiating potential driver mutations from passengers [14]. These various applications of label propagation algorithms highlight its potential in discovering biologically meaningful interactions between a set of genes. Therefore, we used a label propagation-based methodology to connect a given gene set to a larger collection of biological interactions, represented by a network, in order to obtain a more cohesive look into the biological state that is represented by this set of genes and reveal the signaling events that connect the genes to each other and to the broader molecular machinery.

## Results

In this study, we used a label propagation-based methodology to create networks of signaling changes that are observed downstream of common oncogenic genomic alterations across different tumor types, with a particular emphasis on recurrent mutations. Gene expression signatures, consisting of genes differentially expressed in the presence of a given mutation, were used as seeds in a label propagation algorithm to explore a network of known signaling and regulatory pathways. The resulting network for each individual mutation represents the range of molecular events that are dysregulated, revealing the specific signaling and regulatory pathways that are perturbed as a result of this particular genomic alteration. Comparing networks of different mutations highlighted similarities in signaling pathways that were observed downstream of distinct mutations, revealing previously unappreciated connections between the genes that drive cancer and signaling events tumors converge on through distinct mechanisms.

### Label propagation creates networks from gene sets

Label propagation based approaches have the potential to fill in the missing links between a set of genes based on the connectivity information provided by a network. In this study, label propagation was used to create networks of signaling and regulatory pathways starting from a set of genes representing a biological state. This set – which will also be referred to as seed gene set – can be any set of genes one is interested in analyzing in more detail, such as gene expression signatures predicting patient prognosis, a group of genes correlating with drug response, or a set of differentially expressed genes that can discriminate one phenotypic group from another. The second required input is a graph of biological interactions, where nodes represents genes and edges represent functional relationships between them. This network can be generated based on a variety of sources, including results of high-throughput experiments or literature curation. In this particular instance, we chose NCI’s Pathway Interaction Database (PID) [15] as our source of network, mainly because PID is a high-quality resource curated by experts that represents a relatively comprehensive set of currently known signaling and regulatory pathways [16]. After mapping the seed gene set to this network, an iterative diffusion process starts from seed genes, spreading information to the neighboring genes following the paths imposed by the network’s structure. The end result is a subnetwork that contains seed gene set and the additional genes that are reached through the propagation process. The initial gene set will typically be a collection of genes sparsely representing a biological state and this subnetwork that is expanded from them fills in the blanks and gives us a more comprehensive look into the molecular events represented by the gene set. Studying these individual networks in detail highlights the range of pathways that is represented by the initial gene set.

Additionally, we use a distance metric based on the idea of maximal common subgraph [17] to compute pairwise distance between different networks. Smaller distances and higher similarity observed between networks reveals the set of networks that contain overlapping molecular events, which helps us discover unexpected connections between different gene sets, whereas high distances between networks reflect the sets of genes that represent distinct signaling pathways. Figure 1 provides a conceptual summary of this workflow and implementation details are provided in Methods section.

**Figure 1:**
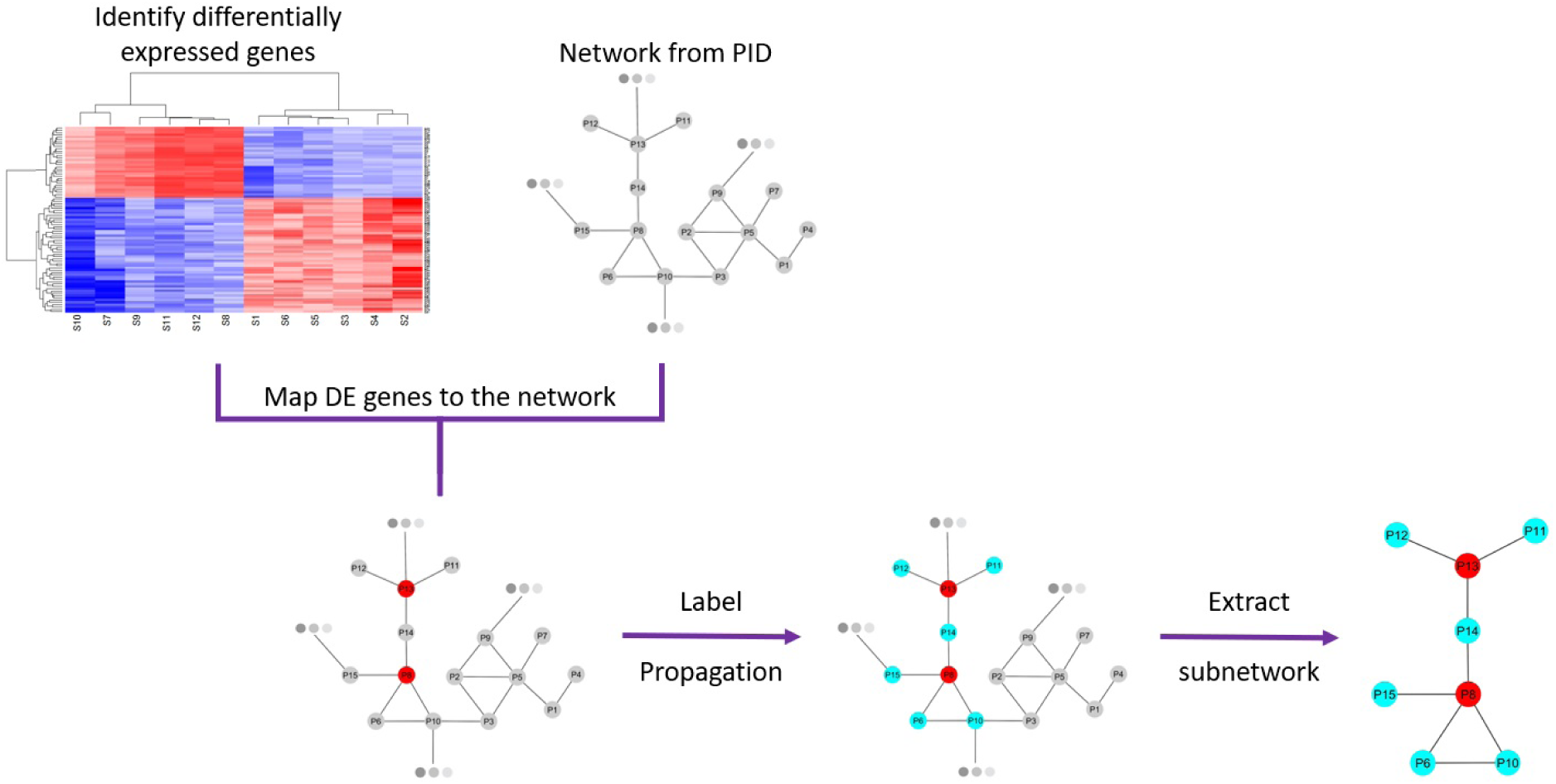
Description of the workflow. The analysis starts with identifying a set of differentially expressed genes (or any gene signature of interest) and obtaining a network of known biological interactions. Label propagation is run starting from the nodes corresponding to the gene signature, in the end to obtain a subnetwork of interactions connecting these genes to additional functionally related genes revealing signaling events represented by the gene set.

### Label propagation recovers pathways

After establishing the workflow, we used a variety of seed gene sets to assess if this approach can successfully connect a particular gene set to the relevant biological pathways that it represents and generate a more comprehensive view of the molecular state that is sparsely represented by the gene set. The first case focused on a controlled set of seed genes, in order to test if the algorithm is capable of identifying multiple distinct pathways when the seed gene set contains genes representing a mixture of pathways. To form this set, we picked four individual pathways that have roles in DNA damage response and repair – namely “Fanconi anemia pathway” [18], “ATR signaling pathway” [19], “ATM pathway” [19] and “p53 pathway” [20]. Five genes were randomly selected from each pathway to create a seed gene set containing 20 genes. The subnetwork obtained at the end of this label propagation run is shown in Figure 2a. In this network, seed genes are color coded based on the pathways to which they belong along with the edges that are identified to be parts of these pathways. As can be seen, when starting with a gene set containing elements from multiple different pathways, the algorithm can recover parts of each individual pathway. A gene expression signature will typically contain components of numerous pathways, therefore it is important to see that the algorithm is capable of diffusing to additional participants of multiple distinct pathways simultaneously. Closer look into the complete set of genes and pathways present in this subnetwork reveals additional interesting biological connections. In addition to the four pathways making up the seed gene set, the network in Figure 2a contains “BARD1 signaling events” pathway, which is also known to have a role in DNA damage response [21]. This means that the algorithm not only recovers missing parts of separate pathways represented by the seed gene set but also links these pathways to additional related pathways with which they interact, enabling a more holistic visualization of the molecular mechanisms that are sparsely represented by the starting gene set.

**Figure 2:**
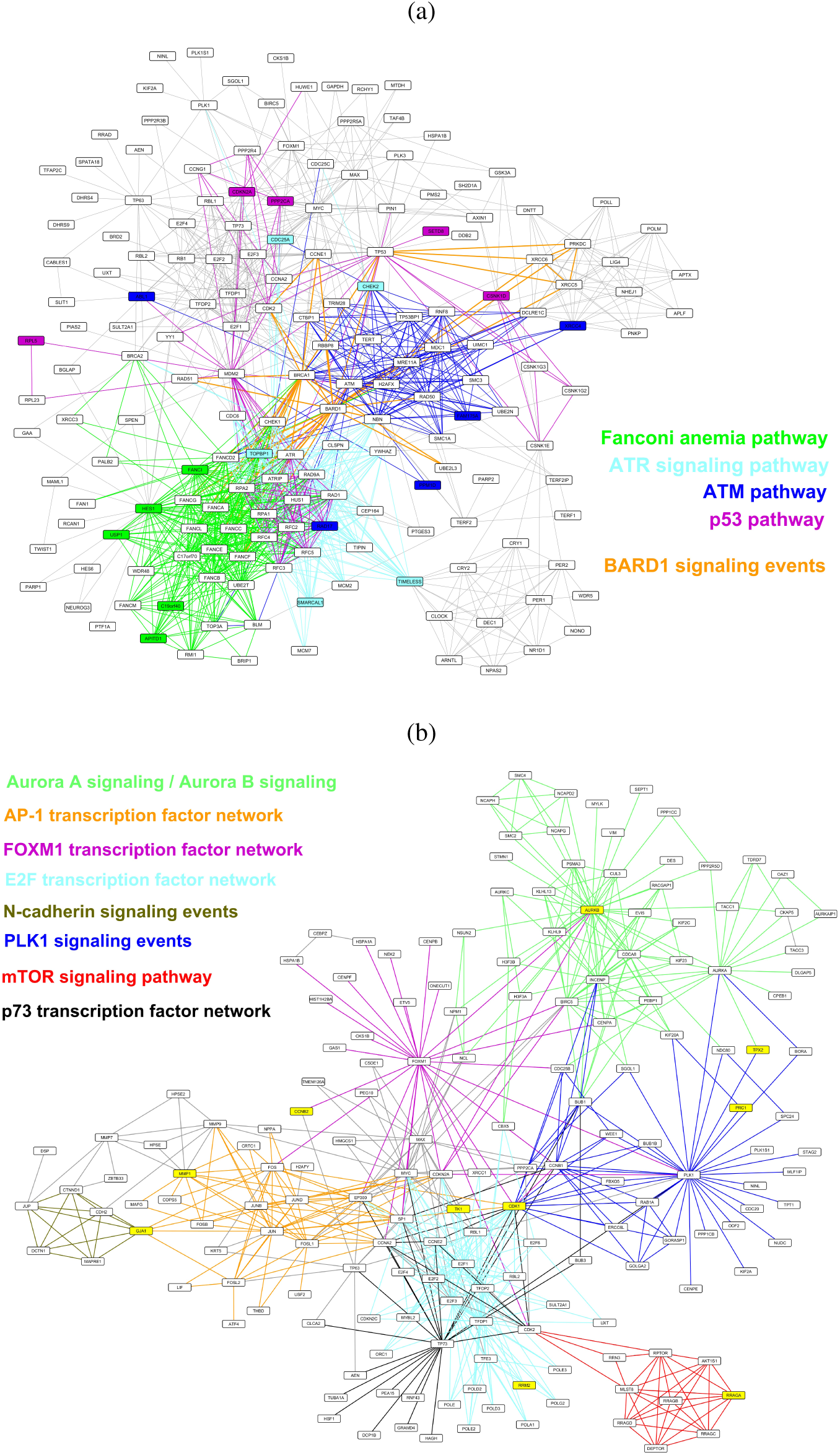
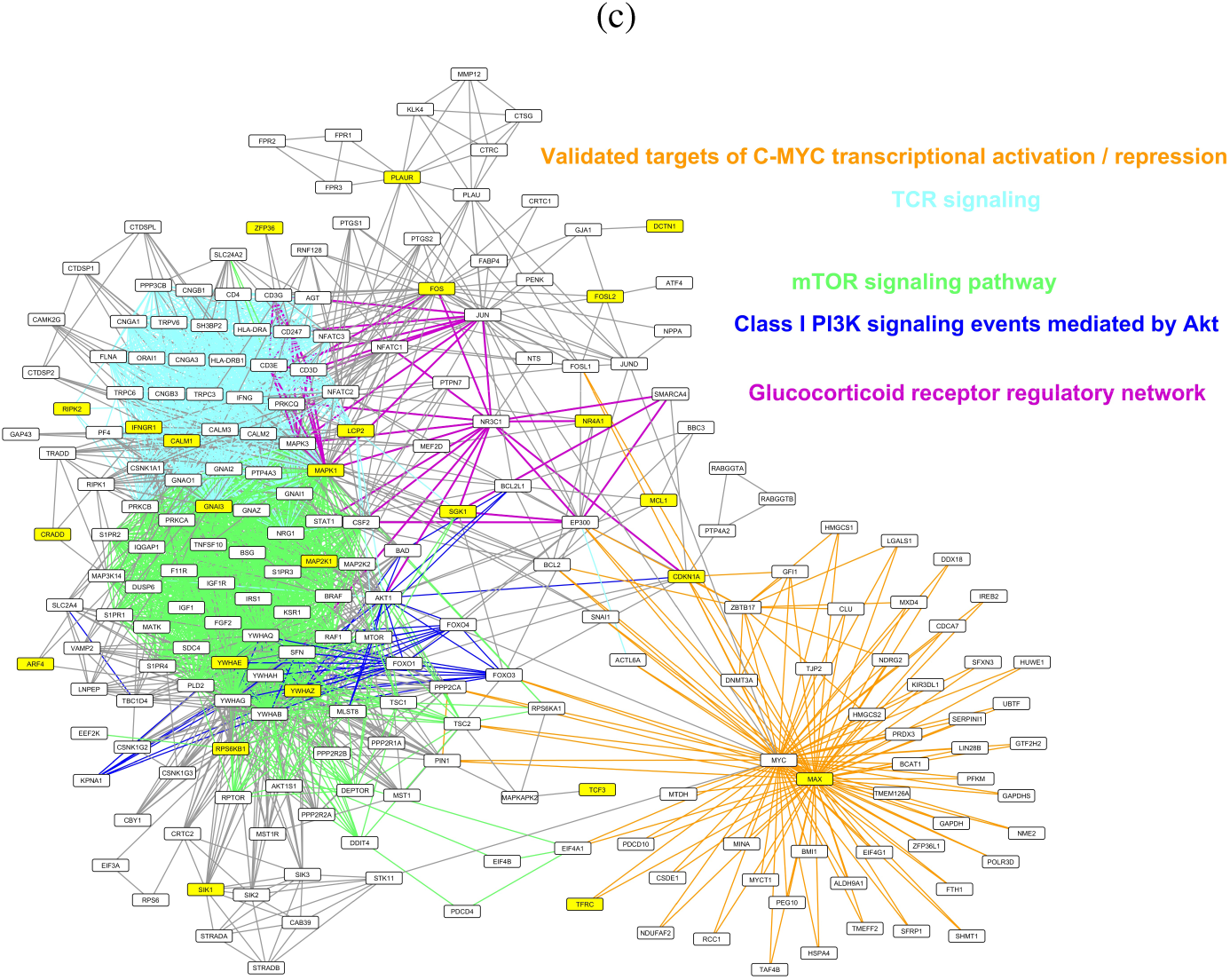
A variety of seed gene sets were used to characterize the workflow’s performance. In all three cases, edges belonging to different signaling events are color-coded and corresponding pathway names are listed next to the networks. (a) A set of genes belonging to four different pathways were randomly selected. The analysis filled in the missing parts of these pathways and additionally highlighted functionally related signaling events. Seed genes are color-coded based on the pathways that they belong. (b) Genes from a “tamoxifen recurrence” signature were used as seed genes to identify the specific signaling events each gene takes part in, highlighting how this approach can be used to improve functional annotations of a given set of genes. Genes from the signature are shown in yellow. (c) Genes from a “glucocorticoid resistance” signature were used as seed genes to identify signaling pathways that have the potential to contribute to the resistance phenotype. Genes from the signature are shown in yellow.

This first gene set was formed by randomly selecting genes from a group of pathways and every pathway had equal number of representatives. However, in most applications of this workflow, the participants of the seed gene set will be dictated by the biological phenomenon that one aims to summarize through a signature and it will be a mixture of pathways, with uneven number of genes belonging to each. To test the performance of the workflow in recapitulating signaling pathways that are represented by a gene expression signature with this make-up, we used a signature that predicts recurrence in breast cancer patients following treatment with the selective estrogen receptor modulator (SERM) tamoxifen [22]. This “tamoxifen signature” contains 36 genes, 10 of which are found in the PID network. These 10 genes were used as the seed genes to run the label propagation algorithm, and the resulting network is visualized in Figure 2b. The edges are color-coded based on the pathways that they participate in, representing the variety of pathways identified. Table 1 provides a more detailed look into the functional roles of these genes based on the annotations provided by Chanrion *et al*. (listed in the middle column) and the corresponding signaling pathways we identified through label propagation. As it can be seen from this table, there is a substantial agreement between these two annotations. For instance, 5 out of these 10 genes have roles in mitosis and cell cycle and they all are represented here as parts of pathways that have well-established roles in mitotic machinery or cell cycle regulation, such as “Aurora A / B signaling” [23] [24], “FOXM1 transcription factor network” [25] and “PLK1 signaling events” [26]. *RRM2* and *TK1* are annotated to have roles in DNA repair and replication, respectively, and they lead the propagation to “E2F transcription factor network”, which is known to have roles in both of these processes [27]. *RRAGA*, the only gene in the set annotated to have a role in signaling, is a part of “mTOR signaling pathway”. All these findings highlight that the label propagation approach was successful in linking these genes to biological pathways that are in line with their functional annotations and in discovering meaningful functional connections between these ten genes by localizing them to the pathways that they participate in. As a result, we can go from a list of genes with broad functional roles to a network that positions each gene into a relevant signaling pathway, creating a more informative result that help us better understand each gene’s contribution to the recurrence of breast cancer.

**Table 1:** Tamoxifen signature. First column lists genes that are both part of the signature and PID network. Second column lists annotations of their functional roles, determined by [22]. Third column lists pathways each gene belongs to in the network shown in Figure 2b.

| Gene | Functional Role | Pathway in PID |
| --- | --- | --- |
| GJA1 | Cell growth, apoptosis, adhesion | AP-1 TF network;<br>N-cadherin signaling events |
| RRM2 | DNA repair | E2F TF network |
| TK1 | DNA replication | E2F TF network;<br>Validated targets of C-MYC transcriptional activation |
| MMP1 | Invasion | AP-1 TF network; Syndecan-1-mediated signaling events |
| AURKB | Mitosis, cell cycle | Aurora A/B/C signaling; FOXM1 TF network |
| CCNB2 | Mitosis, cell cycle | FOXM1 TF network;<br>Validated transcriptional targets of deltaNp63 isoforms |
| CDK1 | Mitosis, cell cycle | AP-1 TF network; E2F TF network; FOXM1 TF network;<br>PLK1 signaling events; p73 TF network |
| PRC1 | Mitosis, cell cycle | PLK1 signaling events |
| TPX2 | Mitosis, cell cycle | Aurora A signaling; PLK1 signaling events |
| RRAGA | Signaling | mTOR signaling pathway |

Through these two signatures, we observed how label propagation can enable the identification of signaling and regulatory pathways that the elements of a given gene set belong to and highlight the cross-talk between these participating pathways. We now highlight how the more comprehensive look generated by this workflow offers additional insight into the biology summarized by a gene signature that may not be captured when solely focusing on the gene set itself. We used a gene signature generated by Wei *et al.*, which represents correlates of resistance to glucocorticoid induced apoptosis in acute lymphoblastic leukemia (ALL) [28]. Genes in this signature were mapped to the PID network to be used as the seed genes and the resulting network is shown in Figure 2c. Using the Connectivity Map (CMAP) [29] and Gene Set Enrichment Analysis (GSEA) [8], the authors suggested that the PI3K / Akt / mTOR signaling axis has a role in this resistance. Replicating the original study’s findings, the network contains genes and interactions belonging to “mTOR signaling pathway” and “Class I PI3K signaling events mediated by Akt”. The network, however, also contains genes that belong to additional signaling pathways not highlighted by the study, and a more detailed look into the individual pathways represented underscores the possibility of these events contributing to the resistance. For example, “TCR signaling” is one of these pathways, and studies by Jamieson et *al*. [30] and Ko et *al*. [31] revealed a role for T-cell receptor signaling in preventing glucocorticoid induced apoptosis. “Validated targets of C-MYC transcriptional activation” and “Validated targets of C-MYC transcriptional repression” are two other pathways identified in this network, and multiple studies have defined a link between the suppression of MYC expression and glucocorticoid induced apoptosis, validating the cross-talk between these pathways [32] [33] [34]. Additionally, a study by Da Costa et *al*. proposed treatment with the BET bromodomain inhibitor JQ1, an inhibitor of MYC transcription, as a way to sensitize ALL cells to dexamethasone treatment [35]. All in all, this label propagation-based network that offers a more thorough understanding of the individual signaling pathways represented by the signature revealed the presence of additional molecular events that have the potential to contribute to resistance. Based on the three cases shown in Figure 2, we can conclude that the workflow is effective in converting a gene set to a signaling network composed of interactions corresponding to their functional roles, generating a comprehensive look into the molecular events represented by the gene set.

### Tumors converge on select signaling pathways downstream of distinct mutations

Thanks to large scale sequencing efforts like The Cancer Genome Atlas (TCGA) project, we have the opportunity to take an in-depth look into the genomes of a wide variety of tumor types. Studies that focus on analyzing the breadth of genomic alterations observed across tumors revealed that the landscape of mutations in different tumor types is variable [4] [5]. Certain genes, like *PIK3CA* and *TP53*, are mutated at a high frequency across a number of different tumor types, whereas there are other genes whose mutations are only observed in a single tumor type. Additionally, different tumors select for distinct sets of mutations to support their genesis and progression, leading to unique mutational profiles for each tumor type. Analyses that focus on uncovering downstream effects of mutations can reveal signaling and regulatory pathways that are dysregulated as a result of these genomic alterations, through which we can gain additional insight into how these mutations contribute to tumor development and progression. Furthermore, performing comparisons across different mutations have the potential to highlight common molecular events that occur downstream of different mutations, revealing recurring signaling changes that different tumor types frequently exploit through distinct genomic alterations. To explore these possibilities, we decided to take a closer look into the sets of frequently mutated genes across a range of tumor types. We picked three different tumor types that have at least ten different genes frequently mutated in the TCGA sample set – urothelial bladder carcinoma, lung adenocarcinoma, and endometrial carcinoma. The following sections focus on the analysis of mutations observed in these tumor types with the label propagation approach and how we used pairwise distances between networks to highlight signaling events tumors converge on through genomic alterations in distinct genes.

#### Networks of mutations observed in bladder carcinoma

Genes frequently altered in bladder carcinoma patients affect crucial cellular processes, such as kinase signaling, histone modifications or cell cycle progression and for some of these genes, such as *CDKN1A* and *RXRA*, bladder cancer is the first tissue type within TCGA studies where they have been observed to be significantly mutated [1]. With analysis detailed below, we explored how some of these unique alterations contribute to the dysregulation of key signaling events. As the first step of this analysis, we curated a list of genes that are mutated in approximately at least 10% of bladder carcinoma patients enrolled in TCGA study, based on the sets of genes that are identified to have frequent mutations in bladder cancer by TCGA study [1], the study by Kandoth *et al*. [4], and TumorPortal [5] (Figure 3a). For each gene within this list, we created an individual signaling subnetwork based on the workflow detailed in Figure 1. A gene signature composed of genes differentially expressed in the presence of a given mutation formed the seed gene set for label propagation. As a result, we obtained a set of networks revealing the underlying biology represented by the dysregulated genes, which in turn reveals the downstream effects of selected mutations. We were especially interested in the relationships between these networks and how they could reveal common signaling events that are affected by distinct genomic changes. To explore these trends in a quantitative manner, we used the distance metric described in Figure 7 to compute the distances between every pair of network. Through this, we generated a pairwise distance matrix that described the relationships between different mutations. To reveal the patterns of similarity, we performed hierarchical clustering on this matrix and the resulting heatmap is shown in Figure 3b.

**Figure 3:**
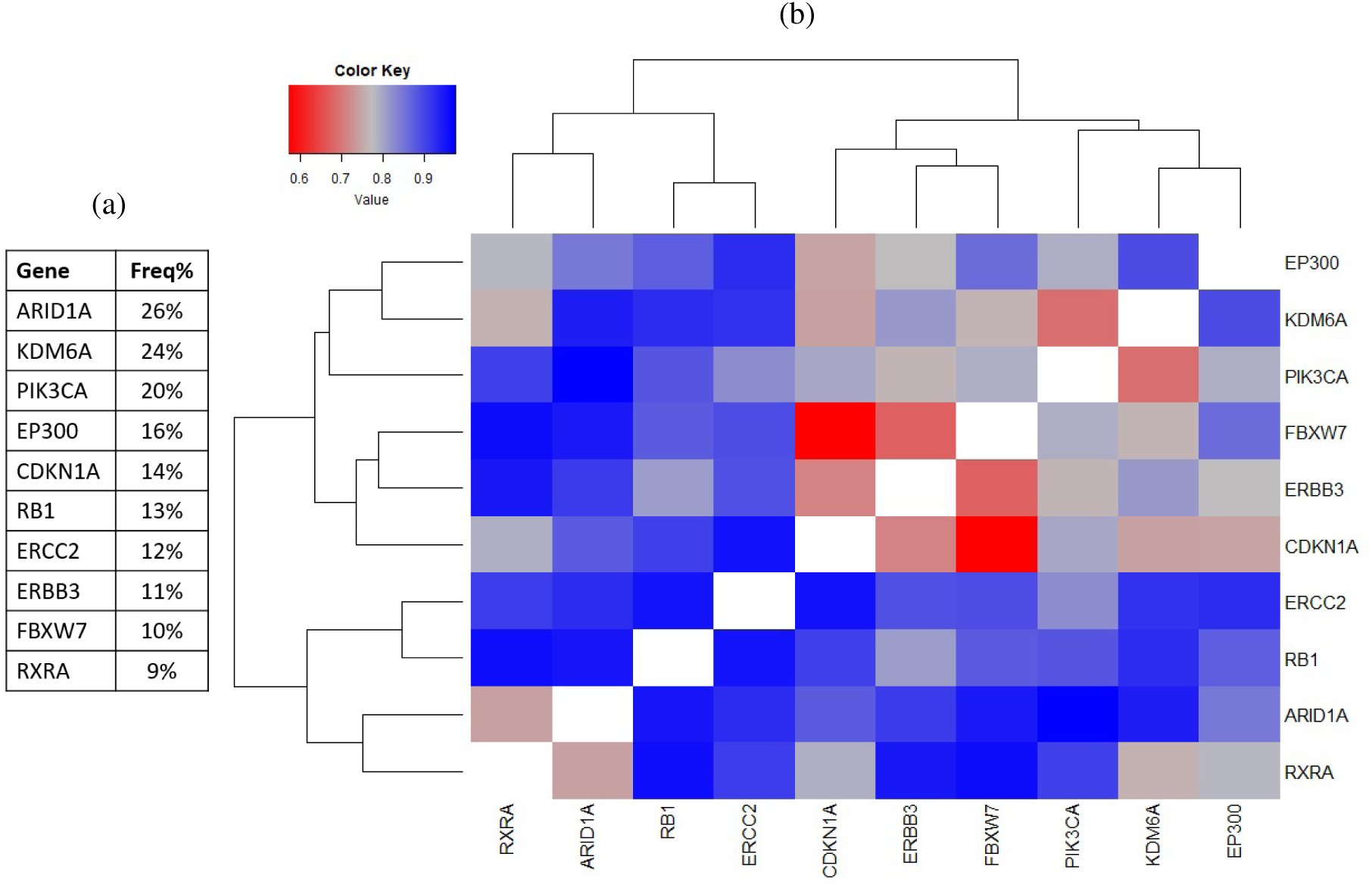
Signaling networks of mutations observed in bladder carcinoma (a) List of genes included in this analysis and their frequency in TCGA population (b) Hierarchical clustering of the matrix representing pairwise distances between label propagation-based networks of genes frequently mutated in bladder cancer. Color key is shown in the upper left corner.

One pattern that this heatmap emphasizes is the fact that within the context of signaling pathways that were analyzed, the majority of networks do not share substantial similarities with others. This implies that distinct genomic alterations have distinct downstream consequences that affect disparate parts of the molecular machinery. The rare cases where we do observe similarity then become more intriguing as these cases have the potential to reveal downstream signaling events that are dysregulated through more than one mutational mechanism. For instance, the most similar pair in bladder carcinoma set is formed by the networks of FBXW7 and CDKN1A. This similarity cannot simply be explained by a high co-occurrence of *FBXW7* and *CDKN1A* mutations as the frequency of patients with mutations in both genes is only 1.5%. We therefore looked further into these two genes to see if there is any overlap between their functional roles that could explain this observed similarity. *FBXW7* encodes for an F-box family member protein, which has roles in substrate recognition of SCF ubiquitin ligases [36]. This ubiquitin ligase has important roles in contributing to cell homeostasis by regulating the proteolysis machinery, and one important target recognized by FBXW7 is Cyclin E [36] [37]. The CDK2 – Cyclin E complex has important roles in regulating cell cycle progression [37] and loss of *FBXW7* activity is expected to lead to the dysregulation of the intricate Cyclin E balance, which in turn will contribute to dysregulation of cell cycle. The other gene in this pairing, *CDKN1A*, encodes for the cyclin dependent kinase (CDK) inhibitor p21 [38]. p21 is a critical regulator of cell cycle through its inhibitory activities on CDK-cyclin complexes, including CDK2 – Cyclin E complex [38] [39]. These established roles of *FBXW7* and *CDKN1A* show that the regulation of CDK2 – Cyclin E complex activity is a common downstream target. Therefore, the similarity we observed in this pair is likely to be driven by the signaling changes that occur upon dysregulation of CDK2 – Cyclin E complex. The pathway view of genes frequently altered in bladder carcinoma generated by TCGA [1] coheres with this observation, as both *CDKN1A* and *FBXW7* are listed as negative regulators of *CCNE1* and cell cycle progression. Through this analysis, we observed that majority of frequently altered genes in bladder carcinoma patients led to minimally overlapping signaling changes, emphasizing the variety of signaling events that are dys-regulated during tumorigenesis. Furthermore, the convergence observed downstream of CDKN1A and FBXW7 networks highlighted the importance of tight regulation of cell cycle progression and multitude of ways tumors disrupt this machinery. Paired with the fact that *CDKN1A* is rarely significantly mutated in other tissue types, this proposes dysregulation of CDK-cyclin complex activity as a critical contributor to the development and progression of bladder cancer.

#### Networks of mutations observed in lung adenocarcinoma

Lung adenocarcinoma is an aggressive disease with high rates of somatic mutations, where a multitude of genes are significantly mutated across the patient population [2]. Some of these genes are well-known and druggable drivers, such as *EGFR*, whereas we don’t have a clear understanding of how others, such as *TSHZ3* and *NAV3*, contribute to lung adenocarcinoma progression. To study the perturbations to signaling pathways downstream of these mutations and the signaling events distinct mutations converge on, we applied the workflow described above to lung adenocarcinoma dataset. Genes selected for this analysis based on the literature [2] [4] [5] were mutated in at least 7% of patients in TCGA study (Figure 4a). For each gene in this list, a network of downstream signaling changes was created as described. Then, pairwise distances between each pair of networks were computed to create the pairwise distance matrix of lung adenocarcinoma. The heatmap obtained after performing hierarchical clustering on this matrix is shown in Figure 4b.

**Figure 4:**
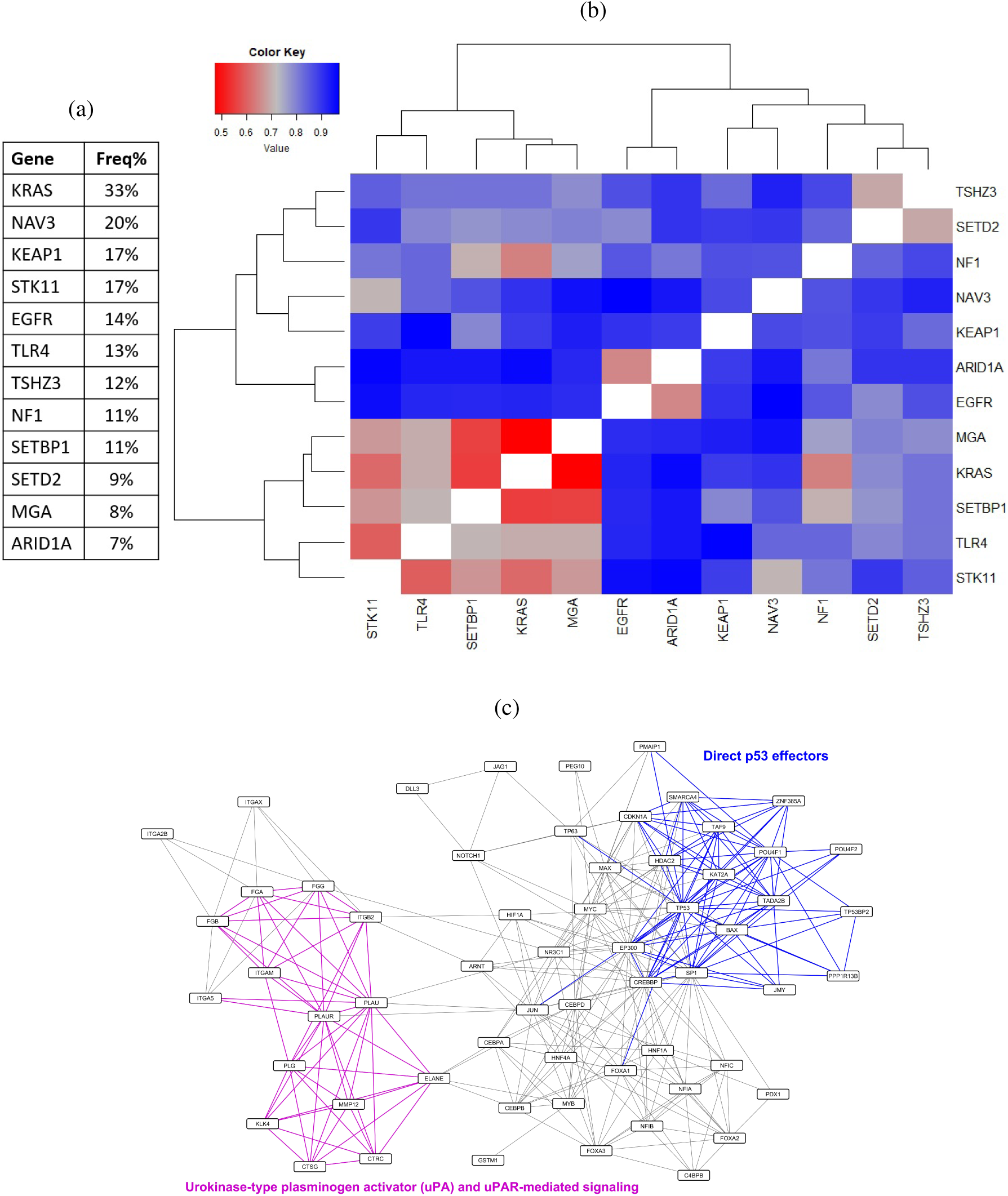
Signaling networks of mutations observed in lung adenocarcinoma (a) List of genes included in this analysis and their frequency in TCGA population (b) Hierarchical clustering of the matrix representing pairwise distances between label propagation-based networks of genes frequently mutated in lung adenocarcinoma. Color key is shown in the upper left corner. (c) Maximal common subgraph of TSHZ3 and SETD2's networks. Edges belonging to two signaling events listed in the network are highlighted in corresponding colors.

A closer look into this heatmap reveals some patterns that are in line with our current understanding of the given genes and others that offer additional insight into the functional roles of these mutations. One case that is a prime example for the former is the similarity observed between networks of KRAS and NF1, two genes whose mutations occur mutually exclusive in lung adenocarcinoma patients [2]. *NF1* encodes for a GTPase-activating protein (GAP) that facilitates the inactivation of Ras proteins [40]. As a result, loss of function mutations in *NF1* lead to the loss of an inactivation mechanism of Ras proteins, leading to constitutively active MAPK signaling. Activating mutations in *KRAS* also leads to an active MAPK signaling axis [2]. Therefore, the signaling changes induced by active MAPK signaling would be in the downstream of both *NF1* and *KRAS* mutations and this would be in line with the observed similarities between the two networks. Another pair that can similarly be expected to have shared signaling changes is KRAS and MGA pairing. *MGA* encodes for a transcription factor that is a *MAX* interacting protein [41] and loss of function mutations in *MGA* are proposed as a mechanism for activating MYC signaling [2]. Similarly, there are multiple studies that place *KRAS* in the upstream of MYC signaling [42] [43]. Taken together, perturbations in MYC activity caused by mutations in *MGA* or *KRAS* are likely to explain the shared signaling changes observed between these two networks. These two pairings provide further support to our idea that studying the relationships between these individual networks reveals functional connections between the corresponding genes and how their mutations can exert shared effects on signaling events, especially highlighting the range of ways a cell can dysregulate an individual pathway.

Following these examples, we turned our attention to pairs of mutations whose overlapping biological functions are less intuitive, in order to see whether we can gain more functional information about a given gene or the pathways that are regulated by it by studying its network and similarity patterns. One such example is *TSHZ3*, which encodes for a zinc finger transcription factor [44]. The network that is the most similar to TSHZ3’s network belongs to SETD2, which encodes for a histone methyltransferase [45]. With this pairing, it is harder to immediately realize a link between the two proteins and their effects on cell signaling, especially because of our limited knowledge on the role of *TSHZ3* in lung cancer. However, analyzing the maximal common subgraph of the two networks revealed clues about common downstream effects of these genes and a potential insight into *TSHZ3*'s contribution to lung adenocarcinoma biology. Figure 4c shows the maximal common subgraph of TSHZ3 and SETD2 networks and one particular pathway of interest – “Direct p53 effectors” – is highlighted in blue. The presence of this signaling event in both of these networks implies that p53 and genes that are in the downstream of p53 are affected by mutations in *TSHZ3* and *SETD2*. Supporting this observation, a study by Xie *et al*. revealed a relationship between *SETD2* and p53 [45], in which *SETD2* can contribute to the regulation of p53 signaling by enhancing its transcriptional activities. Additionally, a recent study discovered a connection between *TSHZ3* and p53, where *TSHZ3* is identified as an inhibitor of p53 activity in lung cancer cell lines [44]. Another major contributor to this maximal common subgraph is “Urokinase-type plasminogen activator (uPA) and uPAR-mediated signaling”, highlighted in purple in Figure 4c. Interestingly, plasminogen activator inhibitor-1 (PAI-1), which is the inhibitor of urokinase-type plasminogen activator (uPA), is a known target of p53 [46] [47] and studies show a cross-talk between p53 and plasminogen activator signaling [48] [49]. Overall, these observations highlight p53 signaling and plasminogen activator pathways as critical signaling events that are perturbed downstream of mutations in *TSHZ3* and *SETD2*, proposing a role for these mutations in lung adenocarcinoma tumorigenesis and progression through regulation of p53’s transcriptional activity. This also shows the value of focusing on the signaling networks and their similarity patterns and how it can reveal underappreciated functional roles of frequently altered genes and ways they contribute to tumor biology.

#### Networks of mutations observed in endometrial carcinoma

High frequeny of mutations in *PTEN* and *PIK3CA* genes implicate PI3-kinase signaling pathway as one of the major drivers of endometrial carcinoma. Additional frequently altered genes have the potential to contribute to dysregulation of this pathway or alternatively perturb other signaling events. To explore the relationships between these alterations, we followed the established workflow using endometrial carcinoma patient samples. Figure 5a shows the selected list of genes that are mutated in at least 10% of endometrial carcinoma patients in TCGA study [3] [4] [5]. Same analysis workflow was used to create individual networks of signaling changes for each gene given in this list. Following that, pairwise distances between these networks were computed and the resulting heatmap when this pairwise distance matrix was clustered is shown in Figure 5b.

**Figure 5:**
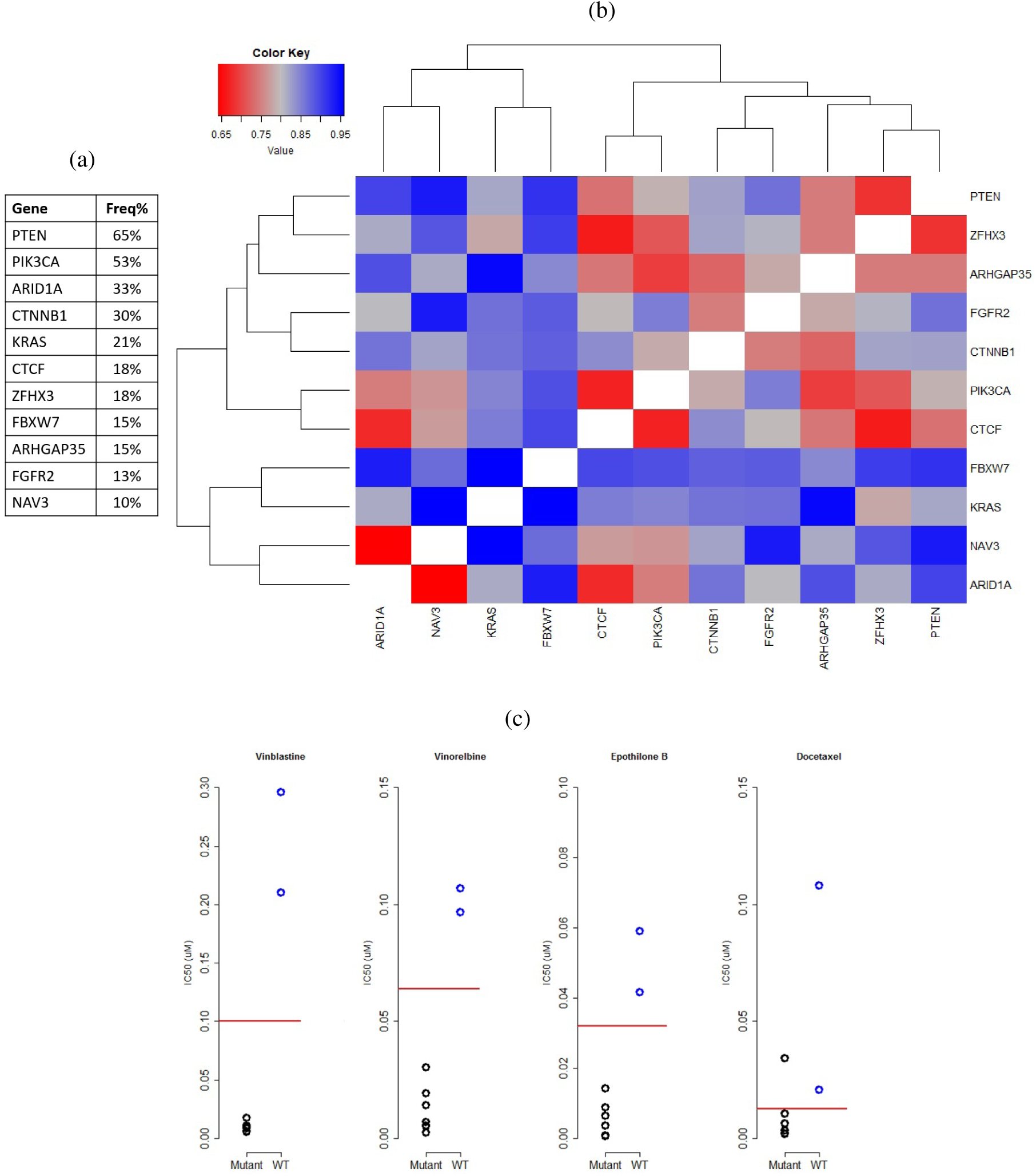
Signaling networks of mutations observed in endometrial carcinoma (a) List of genes included in this analysis and their frequency in TCGA population (b) Hierarchical clustering of the matrix representing pairwise distances between label propagation-based networks of genes frequently mutated in endometrial carcinoma. Color key is shown in the upper left corner. (c) Each graph depicts IC50 values of a microtubule inhibitor measured across a set of endometrial carcinoma cell lines, obtained from GDSC. Black circles represent cell lines that have mutations in either *CTCF* or *ZFHX3* whereas blue circles represents cell lines that are wild-types for both. Red lines represent the maximum screening concentration of each drug.

Loss-of-function mutations in *PTEN* and gain-of-function mutations in *PIK3CA* are both expected to lead to the activation of PI3-kinase signaling pathway [50]. In our analysis, we observed some overlap between these two networks, in line with this expectation. However, there were also signaling events that are uniquely observed downstream of *PTEN* and *PIK3CA* mutations. Studies that looked into the roles of these mutations in detail have highlighted the nonoverlapping roles these two genes have in development and progression of endometrial carcinoma, and molecular events – apart from PI3K pathway activation – that occur downstream of *PTEN* mutations [50] [51]. The unique signaling events observed in PTEN or PIK3CA networks are therefore representative of nonredundant roles of these genes. Additionally, there are several other pairs of genes with higher levels of similarity, such as networks of ARID1A and NAV3, ARID1A and CTCF, or PIK3CA and CTCF. Future work on these pairings have the potential to uncover unexpected connections between their downstream effects and novel insight into their contributions to endometrial carcinoma.

#### Identifying druggable vulnerabilities common to mutations with similar networks

Through pairwise similarity matrices we computed, we were able to identify networks with high-level similarity that shared molecular events that are dysregulated through multiple genomic alterations. We can take a further look into these networks with high similarities to see if they reveal potential druggable vulnerabilities of tumors harboring particular mutations. When a cell is adapting to the presence of a mutation, it leads to changes in the signaling network which can be captured by the networks we created based on dysregulated gene expression patterns. One potential consequence of this rewiring is changes in the drug response profile. Certain mutations can lead to unexpected vulnerabilities in the cell, rendering the tumor sensitive to a certain class of drugs whereas others could affect the molecular architecture in a way to induce resistance to anticancer therapies. Bearing this phenomenon in mind, we can hypothesize that distinct mutations leading to similar changes in the signaling network of a given tumor type have the potential to lead to similar changes in drug response profiles.

To test this hypothesis, we turned our attention to Genomics of Drug Sensitivity in Cancer (GDSC) dataset [52]. GDSC dataset offers baseline sensitivities of 1001 cell lines to 265 drugs, which can be used to perform comparative analysis on sets of cell lines and how they respond to the same set of drugs. We used the pairs of genes from endometrial carcinoma analysis with similar networks for this follow-up, with the aim of searching for druggable vulnerabilities that are shared by two mutations with similar networks. Based on the sequencing information available on GDSC cell lines, we stratified nine available endometrial carcinoma cell lines into two groups: the ones that have mutations in either gene in the pair and the ones that are wild-type for both. The drug response profiles of these groups were then compared to see if there were any drugs that show patterns of differential response. Screening across the available drugs and selected list of pairs of genes revealed a potentially interesting pattern of response to microtubule inhibitors when cell lines were stratified based on mutations in *CTCF* and *ZFHX3*. There were four drugs where only the mutant cell lines were responding to them and all wild-type lines were in non-responder category. These drugs were vinorelbine, epothilone B, vinblastine, and OSU-03012. Interestingly, first three of these four drugs all target microtubules. Stratification of the cell lines based on their mutation status and the corresponding separation of response can be seen in Figure 5c. There is one more microtubule inhibitor in this drug panel – docetaxel – and as it can be seen in Figure 5c, only one of the mutant lines fall into the non-responder category in the case of docetaxel and rest of the cell lines behave similarly to the patterns observed with the other three microtubule inhibitors. Even though the sample sizes for the two groups were small, the consistency across drugs targeting the same molecular event makes this pattern stand out. *CTCF* encodes for an architectural protein that regulates 3D structure of the chromatin by creating topological domains and through this, *CTCF* regulates transcriptional activities of a cell [53]. *ZFHX3* encodes for a transcription factor [54] and sequence variants of this gene have been implicated to have an association with atrial fibrillation [55]. Even though *ZFHX3* is frequently mutated in endometrial carcinoma patients, there is not a clear understanding of the functional implications of these mutations in the context of endometrial cancer. Therefore, any insight we can gain on this gene and its downstream effects will be valuable contributions to our understanding of *ZFHX3* and this analysis nominates a potential connection between *ZFHX3, CTCF* and regulation of microtubule dynamics. Additionally, this trend exemplifies a different approach for identifying potential genomic markers of drug response, especially applicable to cases where more than one genomic alteration trends with patterns of sensitivity. Based on the assumption that certain druggable vulnerabilities are linked to specific changes in the signaling network of a cell, we can focus on pairs of genes whose mutations induce similar signaling changes and look for drug response patterns correlating with these convergent pathways, exploring a unique space of combinations of genomic markers. Overall, this underscores the potential of linking dysregulated signaling events shared across distinct genomic alterations to find druggable vulnerabilities of tumors.

## Discussion

Mutations in genes that are critical in maintaining cellular homeostasis are one of the main events that contribute to tumor development and progression. Understanding the functional contributions of genes to the overall signaling network helps us discover the molecular events that are perturbed when they are mutated and how these perturbations contribute to tumorigenesis. For instance, extensive work on the molecular events downstream of receptors like ErbB family paved the way to improve our understanding of how mutations in these genes are used by tumors to ensure survival [56]. Additionally, discovering functional relationships and cross-talk between different genes revealed that tumors may perturb same signaling events through a multitude of mechanisms; for example mutations in numerous different genes can lead to the activation of MAPK signaling [57]. These are especially valuable findings as redundant ways of achieving the same signaling changes have the potential to affect outcomes of therapeutic interventions [58].

These critical insights gained from individually studying effects of mutations have motivated us to perform the analysis workflow described above on a variety of genes frequently mutated across different tumor types, in order to gain a perspective on the breadth of changes observed across different tumors. We were especially interested in approaching this problem from a signaling network based point of view, rather than simply focusing on lists of differentially expressed genes. Genes which display dysregulated expression in the presence of a mutation offer us a snippet of the signaling changes that occur when a cell is adapting to a particular mutation. Label propagation based methodology described in this study expands this limited look into a more cohesive and comprehensive network, highlighting individual signaling or regulatory events that these differentially expressed genes take part in.

These network views obtained with label propagation gave us the opportunity to study the variety of pathways that are affected by different mutations. More interestingly, studying pairs of networks and their similarities revealed the signaling events tumor cells converge on through mutations in distinct, and sometimes seemingly unrelated genes. By computing a distance metric that focuses on shared signaling events, we were able to discover pairs of genes whose mutations lead to the dysregulation of overlapping molecular events. These observations highlight tumors' ability to make use of distinct mutations to achieve similar perturbations in the signaling network. For instance, the similarity observed between CDKN1A and FBXW7 networks underscores the possibility that cell cycle dysregulation in bladder carcinoma can be achieved by directly inactivating a CDK inhibitor or alternatively, by inactivating a regulatory element. Additional insight gained through this approach includes identifying potential roles for genes which we currently have limited information on how they contribute to tumorigenesis. Contribution of p53 and its downstream signaling events, such as plasminogen activator signaling, to the maximal common subgraph of TSHZ3 and SETD2 networks offers further support to the hypothesis that *TSHZ3’s* role in lung adenocarcinoma include regulation of p53 activity [44]. Overall, comparing and contrasting of dysregulated signaling networks of different mutations offered us a unique look into the intricate molecular changes tumors rely on to survive.

Finally, we can use these patterns of similarity to search for novel therapeutic opportunities. Similarities reflected in molecular states induced by two different mutations also have the potential to be linked to similar drug response patterns. This means that we can use genes with similar downstream signaling changes to define a new search space that stratifies samples based on the combination of these genes rather than simply stratifying based on individual genes. This type of stratification scheme has the potential to uncover unique relationships between a set of genomic markers and drug response, that could otherwise be missed if we were looking for each individual gene separately. Therefore, this detailed look into the dysregulated state of signaling networks can inform discoveries of unique markers of drug sensitivity.

## Methods

### Label propagation algorithm

Zhu *et al*. developed a graph-based label propagation algorithm for solving semi-supervised learning problems using harmonic functions [59]. In an instance of graph based semi-supervised learning, there is a graph *G*, which in total has *n* nodes and e edges. These *n* nodes belong into two groups: there are *u* unlabeled nodes and *l* labeled nodes – each with its own label value *y_i_*. The edge set e is represented with a symmetric weight matrix *W*, where *W_ij_* is nonzero if there is an edge between nodes *i* and *j*. We also define a function *f*. For labeled nodes, this function has the value of their labels *y_i_*. The solution to the learning problem is calculating the value of this function *f* for each unlabeled node. These values can then be used to classify unlabeled nodes into distinct classes represented by labeled nodes, typically by picking a threshold value for *f* and separating unlabeled nodes into classes accordingly by comparing their *f* value with the threshold. The following iterative algorithm can be used to perform these calculations:

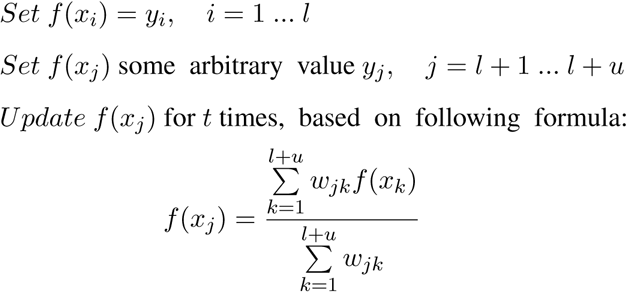

This is the generic description of the label propagation algorithm that we used in our approach. In our context, the graph will be a biological network, where nodes represents genes and edges represents interactions or functional relationships between them. The gene set of interest will form the labeled node set, which means that there will only be one class of labeled nodes. As a result, interpretation of the resulting *f* values will be different than the generic case described above. Rather than picking one threshold value and comparing each unlabeled node’s value against it, we will assess whether an unlabeled node’s *f* value is significantly higher than a value that can be obtained by chance alone. This way we will be focusing on identifying unlabeled nodes which receive a significant amount of diffusion from labeled nodes. To achieve this, we first generated a set of random networks. We used a network randomization algorithm based on “edge switching” principle described by Maslov and Sneppen [60]. To perform randomization through edge switching, first a pair of edges A-B and C-D are selected randomly. Then, edges between these pairs are removed and new edges between A-D and C-B are introduced. Importantly, multiple edges are not allowed between two nodes so if these new edges already exist in the network, this step is not performed and a different pair of edges is selected. Performing this step for many iterations creates a randomized network from the original, while preserving the degree distribution of the original network. After creating random networks, the label propagation algorithm was run with the exact settings as the original network on each of these random networks individually. p-values for each individual unlabeled node’s *f* value were computed by comparing the values obtained with the random networks with the value obtained with the original network. Then, these p-values can be used to determine which unlabeled nodes have significantly high *f* values. Benjamini-Hochberg correction [61] was applied to offer a more conservative control on false discovery rate.

As mentioned above, we also need to specify an input network. There are a variety of options that can be used, each with different focus. Networks that focus on pairwise interactions between proteins that are obtained via high-throughput experiments, such as yeast two-hybrid, would be useful if one is interested in high coverage of potential interactions. Alternatively, manually curated pathway based networks can be used if the focus is more on functional interactions between proteins in the context of signaling and regulatory pathways. We are more interested in the latter – we would like to connect genes to pathways that they are part of rather than simply seeing if there are interactions between them. Additionally, by placing the genes into their respective positions in signaling pathways, we are gaining information about the pathways that are represented by these genes. For these reasons, we decided to use the Pathway Interaction Database (PID) [15] as our network source. PID is a manually curated collection of human signaling and regulatory pathways. To obtain PID pathway data, we used Pathway Commons (PC) database [62].

Overall, the workflow is as follows. The label propagation algorithm requires two inputs: a weight matrix representing a network and a set of labeled nodes. PID data was converted into a binary, symmetric weight matrix W for the first input. If there is a functional interaction between two genes *i* and *j*, the entries *W_ij_* and *W_ji_* are assigned 1. If there is no interaction between *i* and *j*, they are assigned 0. A set of genes of interest, such as a gene expression signature, formed the second input – set of labeled nodes. Every gene is represented by a unique node in PID network so the genes of interest was mapped to their respective nodes to obtain the set of labeled nodes. To initialize *f* function’s values, labeled nodes were assigned 1 and unlabeled nodes were assigned 0. The iterative algorithm was then run *t* times, which was a value specified by the user. Following this, edge switching algorithm was used to generate a set of 1000 random networks. The label propagation algorithm was then run on each of these random networks separately, with the remaining parameters staying exactly the same. p-values based on these label propagation runs were then calculated to identify the nodes that have *f* values significantly higher than what would be expected by chance. The significance thresholds at this step can be adjusted, based on how stringent one wants results to be.

### Selecting parameters of label propagation

As the number of iterations parameter, *t*, is an important determinant in the results the label propagation returns, we sought to determine a value that will result in a high discovery rate across many different cases. To run tests for determining optimal value of number of iterations parameter, we selected a list of pathways that are represented in the PID network and identified the genes that form these pathways. A subset of genes representing a given pathway was randomly selected to form the set of labeled nodes. Multiple sampling sizes were used for each pathway, resulting in labeled node sets containing 10, 20, 35 or 50 genes and multiple sets were sampled for each size. Multiple starting pathways and differing sizes of initial seed gene sets were used to ensure that the value is not optimized based on a single pathway or starting gene set size value, but rather that it reflects a reasonable value to use across a range of settings. Following this random sampling to generate sets of seed genes, label propagation algorithm was run as described above. Number of iterations parameter was changed each time, and total range of values used were *t* = 5, 10, 15, 20, and 25. At the end of an individual run, recovery ratio of genes belonging to the pathway of interest and additional significant genes that are not part of the pathway were computed. “in-pathway” recovery rate corresponds to the ratio of number of significant genes that belong to the pathway of interest over the total number of genes that form the pathway, excluding the genes that are part of the seed set. “out-pathway” recovery rate is the ratio of number of remaining significant genes that are not part of the pathway of interest to the total number of genes in the complete network that are not part of the pathway of interest. These ratios were computed for every single run. Then, average values were obtained for each different value of the number of iterations parameter by computing the mean of in-pathway and out-pathway ratios obtained with a given *t* value across all different pathways and starting gene set sizes. As it can be seen in Figure 6a, *t* = 5 and *t* = 10 were the best performers in maximizing in-pathway recovery rate, with *t* = 10 returning slightly higher in-pathway ratios under some conditions. As a result, in all upcoming runs, we used 10 as the value of the number of iterations parameter.

**Figure 6:**
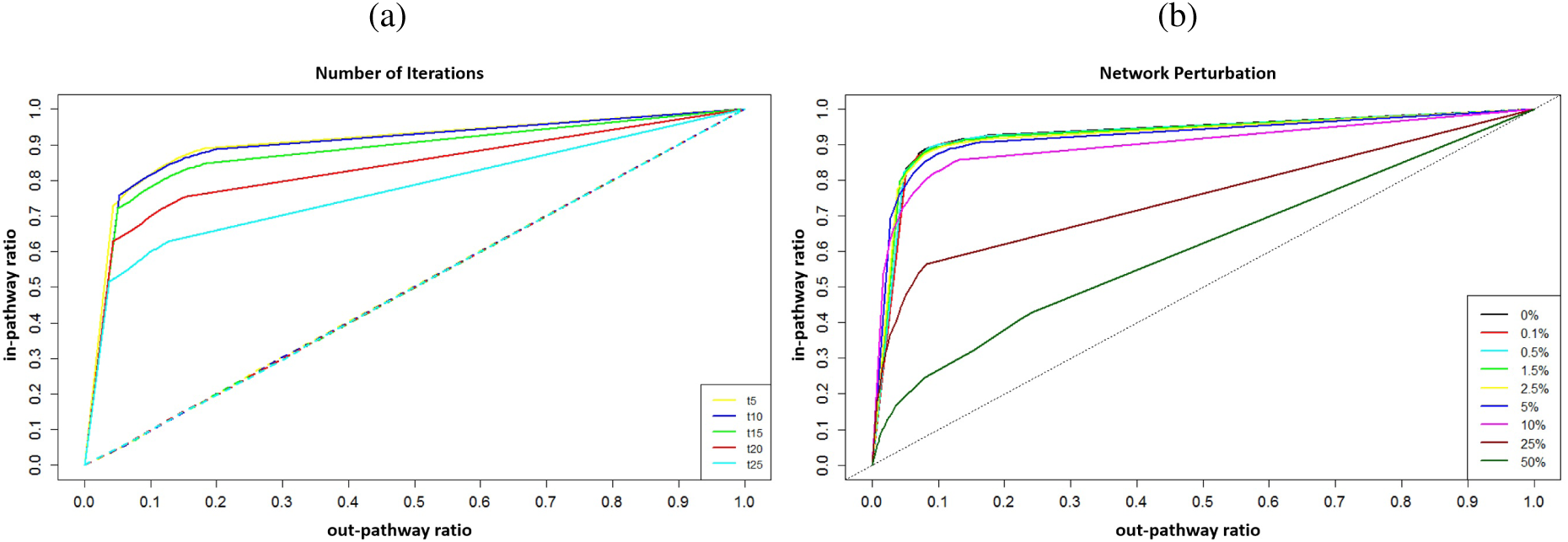
Characterization of the algorithm’s parameters (a) Results of tests performed to identify suitable values for number of iterations parameter (b) Results of tests performed to measure the effects of network structure on label propagation performance

A second round of analysis, shown in Figure 6b, focused on understanding the robustness of this approach with respect to perturbations in the network structure. Specifically, we sought to determine how dependent the results are on the particular topology of the underlying network, which may vary according to human annotation. In a large network like PID, we reasoned that changing the connectivity of a very small percentage of edges should create only minor differences in the complete topology of the network. For instance, the PID network has 33,494 edges, and changing the connectivity of only 34 edges (0.1%) is not a substantial difference in the overall topology of the network. This implies that if we run label propagation on this minimally perturbed network, we would expect to see results that are substantially similar to the results obtained with the unperturbed network. Failure to observe this correspondence would suggest that the subnetworks obtained are not robust to small perturbations and depend instead on a very specific configuration of the complete network. To test this concept, we created a set of networks where a percentage of the PID network’s edges were rewired. To create networks with different levels of perturbation, edge switching algorithm was used with a slight variation [60]. Normally, the number of switching steps performed is a multiple of the total number of edges to ensure the edges are properly mixed. However, in this case, a limited number of switches were performed to ensure that only a subset of edges’ connectivity changes. This threshold was adjusted accordingly to achieve the desired level of perturbation to the overall topology of the network, and at the end, the percentage of rewired edges were 0.1%, 0.5%, 1.5%, 2.5%, 5%, 10%, 25%, and 50%. At each of these levels, multiple perturbed networks were created. For this test, a known pathway from PID was sampled to obtain multiple different sets of labeled nodes. These labeled nodes and perturbed networks were then used as inputs to the label propagation algorithm, where number of iteration parameter was set at *t* = 10. Ratios of in-pathway and out-pathway recovery were computed as described above. Results obtained with networks with same perturbation level were averaged to obtain an individual curve for each perturbation level. Figure 6b shows the in-pathway and out-pathway ratios we obtained at different perturbation levels. The networks with very small perturbation – 0.1% and 0.5% – have curves that are almost the same as the original network, with 1.5% and 2.5% following very closely. After that point, increasing perturbations lead to decreased success in identifying genes belonging to the pathway of interest as the network becomes more and more divergent from the original network. This pattern matches our expectations, which in turn implies that the subnetworks we obtain are robust communities that are connected together through meaningful traversal of the graph and are not spurious subsamplings of the complete network.

### Computing distances between networks

One of the ways we are interested in analyzing the resulting networks is to characterize similarities or differences observed across them. In order to base comparisons between multiple networks on a well-defined quantitative measure, we employed a distance metric that is based on the concept of a maximal common subgraph [17]. This metric, described visually in Figure 7, relies on identifying the maximal common subgraph of two given graphs. When searching for common subgraphs between two graphs, we constrained the search space to the nodes sharing the same label. We imposed this constraint because of the nature of biological networks and information represented by node labels. A biological network's topology represents our knowledge of how genes interact with each other. However, it is not the only source of information in these networks – the labels of nodes are an additional source of information as they represent the genes in a non-redundant fashion. Therefore, when comparing biological networks to identify similarities, it would be misleading to ignore node labels and base the similarity search solely on the connectivity of edges. This might lead to identification of subgraphs with exactly same connectivity but containing different genes and from a signaling perspective, this would represent functionally different subnetworks. Therefore, we instead search for the nodes that are connected in the same manner and that also have the same labels, to make sure that isomorphic subgraphs we identify represents biologically meaningful common subnetworks. Using this constrained search space, we search for a graph *G*_MCS_, where both graphs *G*_1_ and *G*_2_ have a subgraph isomorphic to *G*_MCS_. Of all possible subgraphs that satisfy this criterion, the one with the biggest node size will be the “maximal common subgraph”. Following identification of this subgraph, the distance between two graphs can be computed based on the following formula: 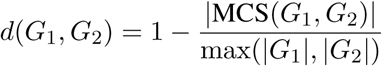

**Figure 7:**
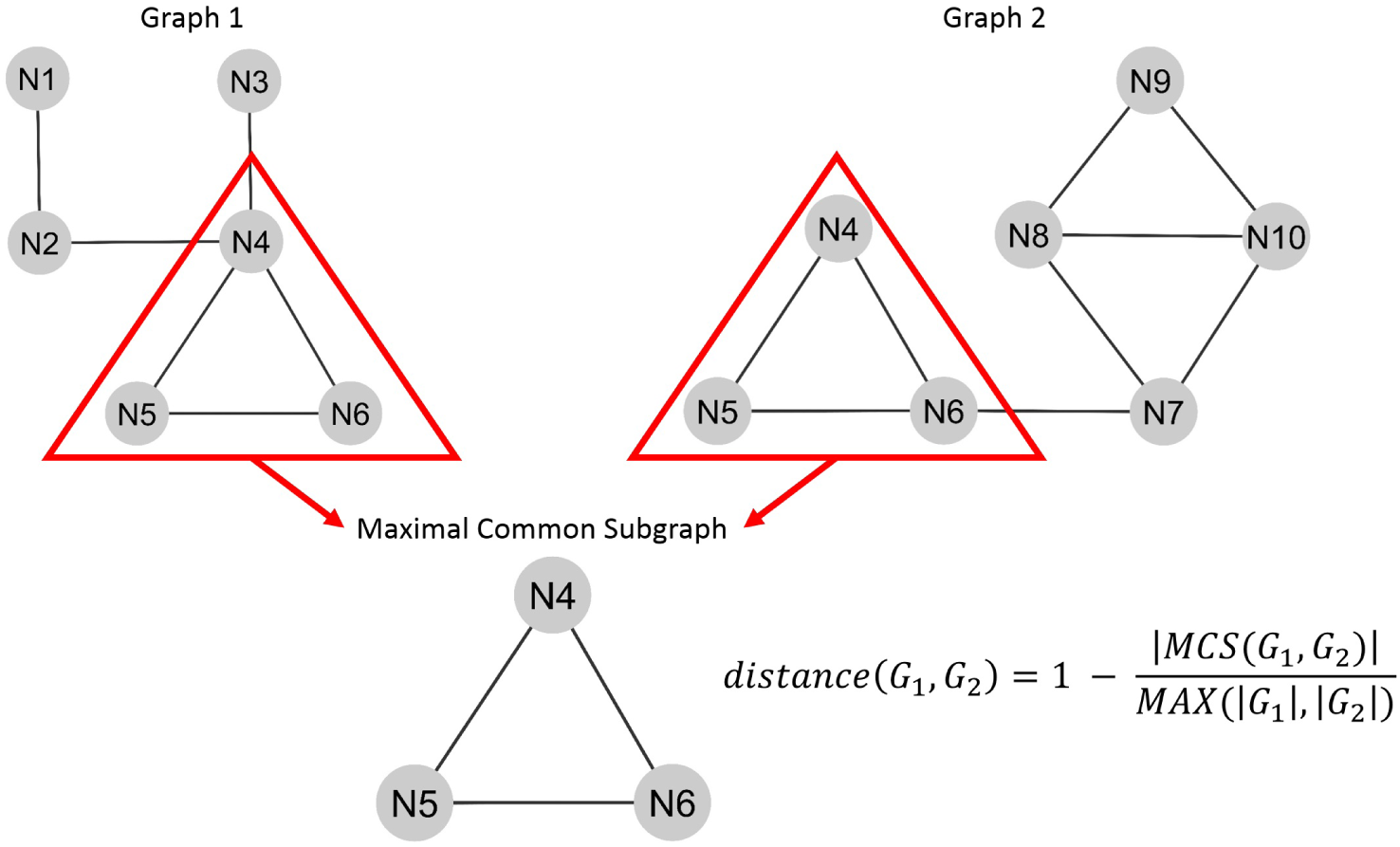
Description of the distance metric. Two graphs are shown to visualize the search space and highlight the corresponding maximal common subgraph.

To test the performance of this distance metric and to assess how well it performs in quantitatively reflecting shared signaling events, we ran a set of tests with pathways found in PID network. We randomly selected five genes from three separate pathways and combined them to form the seed gene sets. “Noncanonical Wnt signaling pathway”, “E2F transcription factor network” and “Regulation of nuclear SMAD2/3 signaling” were the three pathways initially selected for this step. Two separate seed gene sets were created by randomly selecting genes from these three pathways. With each individual set, label propagation was run and then, the distance between these two networks were computed based on the metric described above. This test compares two presumably similar networks, both containing genes belonging to the same three pathways and therefore providing a range of expected distance values that represent similar networks. Following this, we ran additional tests with the aim of creating networks with decreasing levels of similarity to the initial network in order to test if these would correspond to increasing distance values. We generated three sets of seed genes where one of the original three pathways was replaced with another pathway and three sets of seed genes where two of the original three pathways were replaced with other pathways. Finally, to simulate a case with completely different networks, we created two sets of seed genes where the three pathways were completely different from the original three. Particular combinations of pathways and the distance values obtained when these networks were compared to the first network are shown in Table 2. As it can be seen, the distance values are increasing with decreasing level of overlap in the initial set of sampled pathways. The smallest distance is observed when the same set of pathways are used to obtain seed genes. This is followed by cases where two out of three networks stay the same and then by cases where only one out of three networks stays the same. As anticipated, the highest distance values correspond to the cases where seed genes were obtained from completely different pathways. This supports the idea that in cases where we would not expect similarity, the distance metric returns values in agreement with those expectations. Therefore, we relied on this metric to define the levels of similarity observed across different networks when we were searching for convergent signaling events shared across networks.

**Table 2:**
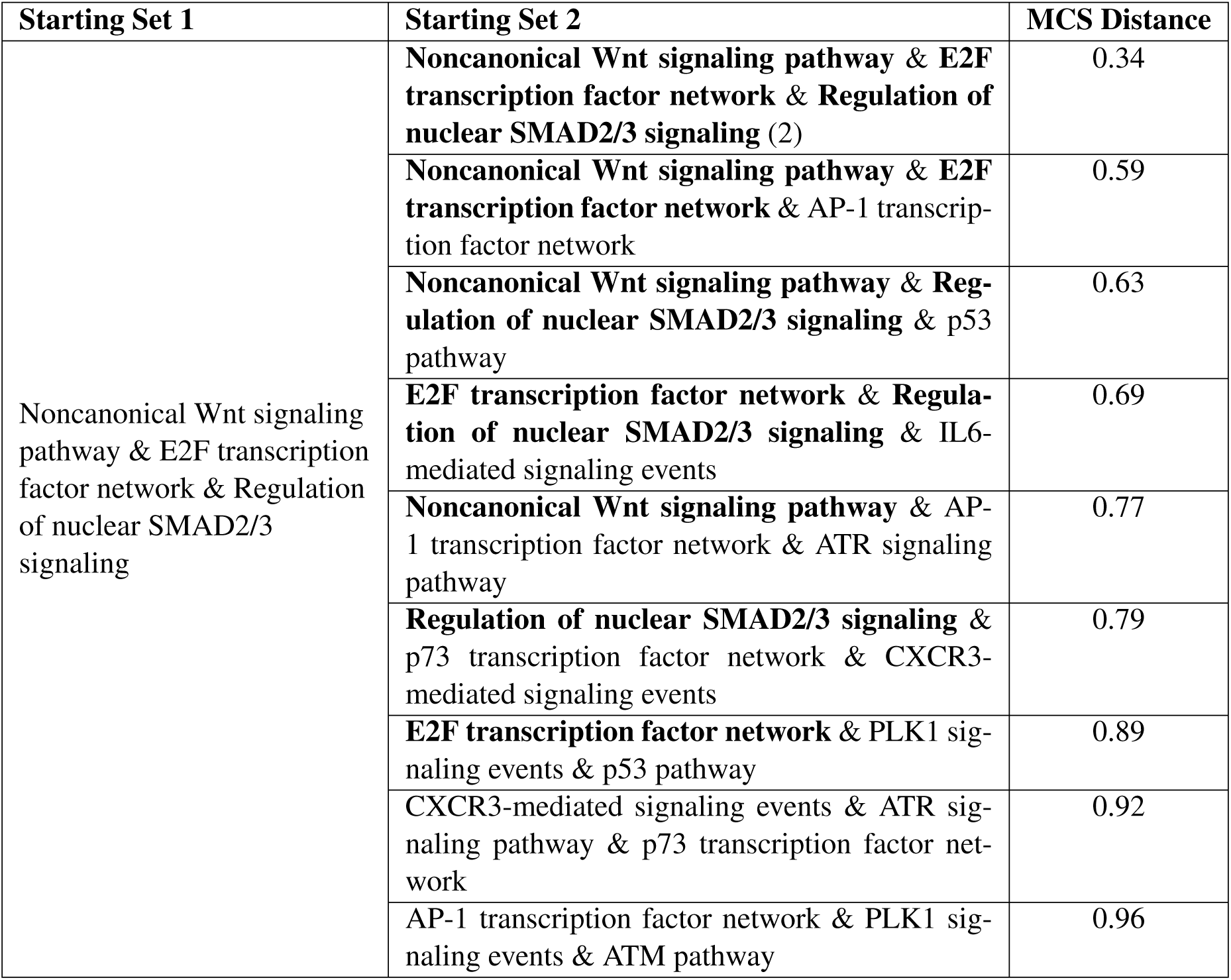
MCS-based distance values obtained across a variety of network comparisons. First two columns list the names of the pathways that contribute to the networks created for these tests. Bold pathway names in the second column represent the pathways that are also part of the “starting set 1” shown in first column. The third column represents the value of distance metric when a given network listed in the second column is compared to the network obtained with starting set 1 seed gene set.

| Starting Set 1 | Starting Set 2 | MCS Distance |
| --- | --- | --- |
| Noncanonical Wnt signaling pathway & E2F transcription factor network & Regulation of nuclear SMAD2/3 signaling | **Noncanonical Wnt signaling pathway & E2F transcription factor network & Regulation of nuclear SMAD2/3 signaling (2)** | 0.34 |
|  | **Noncanonical Wnt signaling pathway & E2F transcription factor network** & AP-1 transcription factor network | 0.59 |
|  | **Noncanonical Wnt signaling pathway & Regulation of nuclear SMAD2/3 signaling** & p53 pathway | 0.63 |
|  | **E2F transcription factor network & Regulation of nuclear SMAD2/3 signaling** & IL6-mediated signaling events | 0.69 |
|  | **Noncanonical Wnt signaling pathway** & AP-1 transcription factor network & ATR signaling pathway | 0.77 |
|  | **Regulation of nuclear SMAD2/3 signaling** & p73 transcription factor network & CXCR3-mediated signaling events | 0.79 |
|  | **E2F transcription factor network** & PLK1 signaling events & p53 pathway | 0.89 |
|  | CXCR3-mediated signaling events & ATR signaling pathway & p73 transcription factor network | 0.92 |
|  | AP-1 transcription factor network & PLK1 signaling events & ATM pathway | 0.96 |

### Analysis of TCGA datasets

RNA-seq gene expression data and mutation data from urothelial bladder carcinoma, lung adenocarcinoma, and uterine corpus endometrial carcinoma patients were obtained through TCGA data portal. For each given tissue, genes of interest were selected based on frequently mutated genes highlighted in their respective TCGA publications, TumorPortal [5], and Kandoth *et al.’s* study across tumor types [4]. For each gene in a given tissue's frequently altered list, we generated a gene expression signature that reflects genes whose transcriptional state is altered in the presence of a particular mutation. For this step, mutation datasets were used to stratify patients into mutant and wild-type groups. Then, gene expression profiles of these two groups were compared to identify genes that were differentially expresed, via Bayesian approximate kernel regression (BAKR) model [63] [64]. This process was performed for each gene individually, in the end to obtain a set of gene expression signatures for each tissue type. These signatures were then used as the labeled node set for label propagation approach, as detailed above. After running label propagation with each signature individually, we obtained a subnetwork of the PID network, representing the signaling and regulatory events that are perturbed downstream of a given mutation. Finally, within each tissue type, distances between all pairs of genes' networks were computed based on maximal common subgraph based metric described above to generate pairwise distance matrices.

### Analysis of drug response

Genomics of Drug Sensitivity in Cancer (GDSC) [52] project was used to obtain drug response profiles and mutation status of screened cell lines. Each drug in this panel was tested across a range of concentrations to create a dose response curve in each cell line and these curves were then used to calculate IC50 values of each drug. For a given drug, we used its maximum screening concentration as a cut-off to separate cell lines into two groups: cell lines with IC50 values greater than maximal screening concentration of a given drug were deemed non-responsive. Remaining cell lines with an IC50 value within the range of screened concentrations formed the responsive group. For each screened drug, panel of cell lines were stratified into these two groups. Additionally, we identified nine cell lines that were categorized as derived from uterine corpus endometrial carcinoma by GDSC and listed pairs of genes with similar networks from endometrial carcinoma analysis, as shown in Figure 5b. For each pair of genes, we stratified cell lines into two groups based on mutation data: those that have mutations in either of these genes and those that are wild-type for both. Our aim was to search for pairs of mutations whose presence placed cell lines into the opposite response category compared to the wild-type cell lines, which can then nominate these genes as candidates for markers of differential drug response profile. For instance, for a given gene pair A and B, we looked for drugs where all responder cell lines had mutations in either A or B and all non-responder cell lines were wild-type (or vice versa). For each gene pairing, we computed the ratio of responder and non-responder cell lines within mutant and wild-type groups separately. Based on these ratios, we identified drugs, where all cell lines in the mutant group were responders (or non-responders) and all cell lines in the wild-type group were non-responders (or responders).

### Network visualization

Cytoscape [65] was used to generate all network views.

